# Financial implications of preterm birth during initial hospitalization: The extent and predictors of catastrophic health expenditure

**DOI:** 10.1101/532697

**Authors:** Hadzri Zainal, Maznah Dahlui, Tin Tin Su

## Abstract

Preterm birth incidence has risen globally and the high cost of initial hospitalization poses financial burden to the family. This study assessed family cost at neonatal intensive care units of two hospitals in the state of Kedah, Malaysia. Family’s expenditure was obtained using a structured questionnaire. 126 families who were government employed spent a mean total cost of MYR 549 (MYR 0 - MYR 4,700) compared to MYR 650 (MYR 40 – MYR 9,300) for 244 families who were not government employed. Mean income loss was MYR 310 (MYR 0 – MYR 15,000) and MYR 348 (MYR 0 – MYR 5,500) respectively. Travel expenses was the cost driver for all families. 15% of families in this study were already living below the income poverty line and majority were not government employed. For the rest of the families, 21% became impoverished when one month household income was used for hospitalization cost but this lowered to 9% with cumulative household income by length of hospital stay. Overall incidence of catastrophic health expenditure among families was 38%. Using multivariable logistic regression household income and residential location were predictive factors for catastrophic health expenditure. Despite universal health coverage through subsidy of direct medical (hospital) cost, the high incidence of catastrophic health expenditure and impoverishment among families of preterm infants was attributable to out of pocket payment for direct non-medical cost (such as travel and food) and indirect cost from income loss. Government employed families with an array of employment benefits appear better protected against financial hardship compared to those in private sector or self-employed. Remedial measures include improving neonatal intensive care unit rooming-in service for mothers, complementary financial assistance for families and enhancing universal health coverage through affordable social health insurance for infant healthcare.

## INTRODUCTION

Preterm birth is defined as delivery before 37 completed weeks of gestation. Preterm birth is increasingly common with substantial medical, economic and social impact as it is invariably associated with acute and chronic complications (1, 2). Since its inception in 2009, Malaysia’s preterm birth registry showed an increasing rate from 8.1% to 11.3% between 2010 and 2012 (3). Due to advancements in care over the last few decades, outcome and survival of preterm infants have improved, however, the economic impact of preterm care has gained much attention (1, 2). Most studies have been devoted to cost of intensive care as initial hospitalization accounted for the bulk of health care cost during the first 2 years of life of a preterm infant (4). However, cost analysis during initial hospitalization had rarely taken into account the family’s perspective (2)

Preterm birth has been found to cause significant out of pocket (OOP) spending for families of preterm infants with estimated cost of up to 2% to 4% of gross annual income during neonatal period (2, 5). For extremely low birth weight infants it was estimated that parental mean cost prior to discharge was up to 4% of the total cost (4). Travel expenses contributed 64% of these non-reimbursable payments, 30% from loss of earnings and 6% from accommodation during visits.

Compounding the situation is the fact that OOP payment is the most prevalent means of healthcare payment in Asian countries and households are at risk of catastrophic health expenditure (CHE) and impoverishment (6, 7). Parents have been advocated as vital providers of informal care or ‘invisible hands’ who should be part of the team caring for preterm infants throughout their hospital stay and this role needs to be acknowledged by formal care providers (8). It is imperative that the issue of financial burden on families especially those with poor socioeconomic background is given due attention by healthcare providers (1). It would ensure a stable home socioeconomic environment that allows parents to focus on providing care and support for infants during initial hospitalization.

## METHODOLOGY

### Study design and participants

This study utilized universal sampling and had prospectively followed up preterm infants from birth till discharge from initial hospitalization. Study was conducted at the neonatal intensive care units (NICU) of Hospital Sultanah Bahiyah (Center 1) and Hospital Sultan Abdul Halim (Center 2), the largest hospitals and referral centres for the state of Kedah which has one of the highest mean number of neonatal admissions per hospital in Malaysia (9). Center 1 offered tertiary level care and had an average of 800 preterm admissions annually while Center 2 had secondary level care with an average of 500 preterm admissions annually. Inborn and out born preterm infants delivered via normal delivery or Caesarean section and admitted to NICU of both centers during data collection period were included in this study. Excluded were preterm infants admitted for less than 24 hours (for observation) and preterm infants with severe congenital anomalies as these infants would only receive supportive care due to their often short duration of life. As incidence of CHE among families of preterm infants during initial hospitalization was not available from literature review, a pilot study for family cost (n=54) was performed over two months (July and August 2015) to provide an estimate that would allow determination of sample size. Sample size calculated with OpenEpi was 370 using pilot study CHE incidence of 40% and 95% confidence interval. Data collection for analysis was conducted from September 2015 to August 2016.

### Cost data collection

Family cost was assessed by a structured questionnaire face to face interview prior to discharge and via phone after discharge (following informed consent). The questionnaire gathered demographic information such as parental age, ethnicity, residential address, education level, occupation and monthly household income. Residential distance from the hospital of admission was categorised as within or outside of the hospital district. Employment of parents in the civil service was important data to obtain as it affected hospital cost payment. Parents were required to provide information on average HHI per month contributed by household members monthly in the form of salary, pension, sale of products, rentals or others. Parents were asked to recall their monthly household expenditure for essentials such as food, housing, utilities, education, transport, healthcare and insurance. Clinical characteristics gathered were gestation, birth weight, delivery mode and length of stay (LOS) of the preterm infant. Both demographic and clinical characteristics were subsequently studied for incidence and association with CHE in families of preterm infants.

Cost incurred by families of preterm infants throughout NICU admission was divided into direct medical, direct non-medical and indirect costs (2). Direct medical cost refers to total hospital cost which families of preterm infants are required to pay upon discharge and commonly OOP payment is the method used. It includes costs of consultation, medication, diagnostic tests and ward fee. Direct medical cost differed between parents who were employed in the public sector and those in the private sector or self-employed. Hospital cost in Malaysia is heavily subsidized by the government with a small contribution from patients as co-payment. However, for parents working in the public sector the cost is fully borne by the government. Those in the private sector, self-employed and non-employed received partial subsidization of hospital cost and are at risk of financial burden due to co-payment. Direct non-medical cost is expenses incurred by parents to enable hospital visitations. It includes transportation, accommodation, meals during lodgings or visits and care of siblings or dependent persons at home. For cost of transportation data was collected on the frequency of hospital visits, transportation mode, distance travelled and expenses for fuel, toll, fares or parking fees paid for each visit. Some families had sought temporary accommodation as they needed to travel for hospital visits. Indirect cost was accounted for by income losses due to altered job schedule and lost wages (if paid by hour or day) and missed working days from time spent for hospital visits that otherwise would have been spent for working. Indirect cost or productive work time lost was assessed using the human capital approach where productive work time lost refers to the work output that would have been generated if the illness event had not occurred (10, 11). The loss of work output is the foregone income of the working individual in the family. Parents were required to convert missed working hours or days into monetary estimates in the questionnaire. This approach was used instead of personnel salary per hour to indicate loss estimation for those with irregular working hours. Income loss was compared in terms of incidence and mean cost between government and non-government employed families.

Total cost per family during initial hospitalization was given by the sum of direct medical cost (hospital fee) and direct non-medical cost (travel, accommodation, food and dependent care). Mean total cost was calculated for government employed or non-government employed families separately. Missed working hours or days and the resultant income loss (indirect cost) for parents due to NICU visitations was categorized as indirect cost. However this was excluded from calculation of total family cost and assessed separately as a measure of financial hardship. Cost items that contributed most to total cost of hospitalization were identified. Costs were described in mean (range) and percentages.

### Financial hardship

The effect of partial (non-government employed parent/s) and full (government employed parent/s) government subsidization of hospital cost on measures of financial hardship was also studied. Hospital (direct medical) cost settlement, perceived difficulty and coping strategies, income loss suffered, CHE and impoverishment were outcomes used as measures of financial hardship. Hospital cost coping strategies and hospital cost settlement were objective assessments while perceived difficulty of paying hospital cost was a subjective assessment of the ability of families to resolve the acute financial burden from hospital admission. Coping strategy methods enquired from parents include use of current income of household member, savings, payment or reimbursement from a health insurance plan, selling assets, borrowing from friends or relatives. Hospital cost settlement looked at whether they were able to make full payment of hospital cost upon discharge or otherwise. Parents were asked in the questionnaire about the level of difficulty experienced paying for hospital cost. Likert scale was used to identify the degree of difficulty ranging from ‘not difficult at all’, ‘not that difficult’, ‘difficult’ and ‘very difficult’. An estimate of income loss from working hours foregone due to hospital visitations (indirect cost) was excluded from calculation of total family cost and assessed separately as a measure of financial burden.

### Impoverishment and Catastrophic Health Expenditure (CHE)

Poverty line is an estimated minimum level of income deemed necessary for basic household needs expenditure in the form of cash, goods and services and impoverishment occurs when households are pushed below or further below the poverty line and other household basic consumption must be forgone (12, 13). Using the 2014 Peninsular Malaysia monthly HHI poverty line of MYR 930 government and non-government employed families of preterm infants were assessed for poverty before and impoverishment after OOP payment of total initial hospitalization (direct medical and direct non-medical) cost (14). In this study impoverishment occurs when total hospitalization cost subtracted from HHI left the family with an income amount below the poverty line of MYR930.

Impoverishment was assessed using two possible scenarios: where one month HHI or cumulative monthly HHI (comparable to LOS in NICU) was utilized. Meanwhile CHE occurs when a household’s OOP payments exceed its available resources and affects the consumption of other necessary goods and services (15). CHE is deemed to occur if OOP healthcare payment exceeds 40% of a household’s capacity to pay (16). A household’s capacity to pay is defined as effective income remaining after basic subsistence needs have been met. Effective income is stated as the total consumption expenditure of the household as an indicator of purchasing power. Effective income remaining is given by total household consumption expenditure minus food expenditure, which equates to non-food essential expenditure as a measure of household capacity to pay.

To determine the incidence of CHE among families of preterm infants, effective household income was first calculated by subtracting food expenditure from total household expenditure. The total cost of hospitalization was obtained by totalling direct medical and direct non-medical costs. CHE was deemed to occur if total hospitalization cost exceeded 40% of effective household income. Total hospitalization cost was the nominator and cumulative effective HHI for the duration of NICU admission was the denominator. For example if a preterm infant’s stay in the NICU exceeded 30 days but was not more than 60 days the cumulative effective HHI used for calculation was 2 months’ worth. Incidence and factors associated with CHE in families of preterm infants were analyzed according to demographic and clinical (gestation, delivery mode, birth weight, length of stay) characteristics.

### Statistical analysis

All cost data were presented in Malaysian Ringgit (MYR) where $1 = MYR 3.75 (prevailing conversation rate in 2015). Cost data was tabulated with Microsoft Excel (2010) and subsequently analyzed with IBM SPSS Statistics version 22 for descriptive and inferential data evaluation. Means with range for continuous variables and frequencies and percentages for categorical variables were used in descriptive analysis. The association between independent factors and the outcome was assessed using univariable and multivariable logistic regression to determine predictors of CHE. Variables were categorised into clinical (preterm infant) and household factors and assessed using univariable logistic regression. Continuous variables were transformed into categorical variables before analysis was performed to avoid outlier effect. The result was interpreted using a P value of <0.05 as a measure of statistical significance. Variables found to be significantly associated with CHE were retained for multivariable regression analysis. Goodness of fit was determined using the Hosmer–Lemeshow test.

## RESULTS

### Respondents’ demography

The response rate among eligible families in this study was 100%. The majority of families of preterm infants were Malays (89%) with Indian (5%), Chinese (4%) and others (2%) making up the ethnic groups (Table 1). 80% of fathers and 90% of mothers were aged between 20-40 years. 1% of fathers and 4% of mothers were below the age of 20. Parents with secondary education were the biggest group (58% respectively for paternal and maternal groups), followed by tertiary education (34% and 36% respectively) and primary level or lower (less than 10% in each group). Only 24% of fathers worked in the government sector while the rest were either self-employed (36%) or in the private sector (40%). Unsurprisingly majority of mothers were homemakers (54%) with the rest in government service (22%), private sector (18%) and self-employed (6%). There were more families of preterm infants who came from outside the hospital district (58%) compared to within the district (42%).

**Table 1:** Respondents’ demographic characteristics (*n*=370)

| Demographic Variables |  | *n* | % |
| --- | --- | --- | --- |
| Ethnicity | Malay | 329 | 89 |
|  | Chinese | 15 | 4 |
|  | Indian | 20 | 5 |
|  | Others | 6 | 2 |
| Paternal Age | <20 | 3 | 1 |
|  | 20-29 | 132 | 36 |
|  | 30-39 | 162 | 44 |
|  | ≥40 | 70 | 19 |
| Maternal age | <20 | 14 | 4 |
|  | 20-29 | 191 | 52 |
|  | 30-39 | 142 | 38 |
|  | ≥40 | 23 | 6 |
| Paternal education | Primary or lower | 26 | 8 |
|  | Secondary | 214 | 58 |
|  | Tertiary | 127 | 34 |
| Maternal education | Primary or lower | 20 | 6 |
|  | Secondary | 215 | 58 |
|  | Tertiary | 135 | 36 |
| Paternal occupation | Self-employed | 135 | 36 |
|  | Government /Pensioner | 85 | 24 |
|  | Private sector | 146 | 40 |
| Maternal occupation | Self-employed | 20 | 6 |
|  | Government / Pensioner | 83 | 22 |
|  | Private sector | 66 | 18 |
|  | Home maker | 201 | 54 |
| Residential distance | Within district | 157 | 42 |
|  | Outside district | 213 | 58 |

Late preterm group comprised the majority of preterm infants recruited (42%) followed by very preterm (28%), moderate preterm (26%) and extreme preterm (4%) groups (Table 2). Most preterm infants recruited had birth weight in the range of 1.50kg to 1.99kg (39%) with almost equal proportions for those in between 1.00kg to 1.49kg (27%) and 2.00 kg to 2.99kg (28%) while the smallest group was in the 0.60kg to 0.99kg range (6%). Majority of them were delivered via Caesarean section (61%). About two-thirds (66%) stayed in NICU for less than a month while close to a tenth (9%) stayed for more than two months.

**Table 2:** Respondents’ child (preterm) admission characteristics (*n*=370)

| Clinical Variables |  | *n* | % |
| --- | --- | --- | --- |
| Gestation | Late preterm | 156 | 42 |
|  | Moderate preterm | 96 | 26 |
|  | Very preterm | 103 | 28 |
|  | Extreme preterm | 15 | 4 |
| Birth weight (kg) | 2.00 - 2.99 | 105 | 28 |
|  | 1.50 - 1.99 | 145 | 39 |
|  | 1.00 -1.49 | 99 | 27 |
|  | 0.60 - 0.99 | 21 | 6 |
| Mode of delivery | Normal | 145 | 39 |
|  | Caesarean | 225 | 61 |
| Length of stay (days) | <30 | 243 | 66 |
|  | 30-59 | 93 | 25 |
|  | ≥60 | 34 | 9 |

### Cost analysis

For families who were government employed the mean total cost during initial hospitalization was MYR 549 (MYR 0 - MYR 4,700) (Table 3). Travel and food expenses contributed 65% and 28%, respectively of total cost incurred during this period. Mean opportunity cost was MYR 310 (MYR 0 – MYR 15,000). Meanwhile for families who were not government employed the mean total cost during initial hospitalization was MYR 650 (MYR 40 – MYR 9,300). Travel accounted for 49% while hospital bills were 37% of the total cost. Mean opportunity cost was higher in this group at MYR 348 (MYR 0 – MYR 5,500). Up to 59% of them incurred income loss compared to 19% of government employed families.

**Table 3:** Family cost for initial hospitalization (*n*=370)

| Cost<br>Component | Cost<br>Item | Families of preterm infants employment |  |  |  |
| --- | --- | --- | --- | --- | --- |
|  |  | Government |  | Non-government |  |
|  |  | Mean (range)<br>MYR | % | Mean (range) MYR | % |
| **Direct medical** | Hospital | full subsidy |  | 242 (0-3161) | 37.2% |
| **Direct non-medical** | Travel | 357 (0-3030) | 65.0% | 315 (0-7800) | 48.5% |
| **Direct non-medical** | Accommodation | 22 (0-1500) | 4.0% | 3 (0-400) | 0.5% |
| **Direct non-medical** | Food | 151 (0-1800) | 27.5% | 84 (0-1635) | 12.9% |
| **Direct non-medical** | Childcare | 17 (0-600) | 3.1% | 5 (0-1000) | 0.8% |
|  | Total | 549 (0-4700) | 100% | 650 (40-9300) | 100% |
| **Opportunity cost** |  |  |  |  |  |
|  | Mean income loss | 310 (0-15,000) |  | 348 (0-5,500) |  |
| | Number of families ($n$) | 126 | | 244 | |
|  | Number incurred | 24 (19%) |  | 144 (59%) |  |

### Financial burden

35% of families of preterm infants in this study received full government subsidy for hospital cost. 25% of families of preterm infants utilized current income of any household member, 18% used savings and 15% resorted to borrowing from friends or relatives to finance the hospitalization cost. 7% of families used combinations of the aforementioned strategies to finance for hospital cost. None of the families had an insurance scheme that provides coverage for newborn infants. 45% of the families with co-payment rated it as ‘not that difficult’ while 25% rated it as ‘not difficult at all’. However 21% of families rated co-payment as ‘difficult’ and 9% rated it as ‘very difficult’. Up to 18% of families with co-payment could not make full payment upon discharge.

Table 4 shows the impoverishing effect of OOP spending (direct medical and direct non-medical costs) for families of preterm infants during initial hospitalization. Overall 55 of 370 families (15%) included in this study were living below the income poverty line and majority of them (96%) were not government employed. Among government employed families only 2 (2%) were living below income poverty line compared to 53 (22%) of those not government employed. Of the 315 families living above poverty line 65 (21%) became impoverished when one month HHI was used for costs during initial hospitalization. When cumulative HHI by LOS was used this figure lowered to 29 families (9%). 6 of 124 (5%) government employed families who were above poverty line became impoverished when one month HHI was used for costs during initial hospitalization compared to 59 of 191 (31%) in non-government employed families. However figures improved when cumulative HHI by LOS was used where only 1 of 124 government employed families who were above poverty line became impoverished and 28 of 191 (15%) non-government employed families.

**Table 4:** Impoverishing effect of initial hospitalization

| Poverty* before initial hospitalization |  |  |  |  |  |  |
| --- | --- | --- | --- | --- | --- | --- |
|  | All families<br>(n=370) |  | Government employed<br>(n=126) |  | Non-government employed<br>(n=244) |  |
|  | Poverty | % | Poverty | % | Poverty | % |
| HHI per month | 55 | 15 | 2 | 2 | 53 | 22 |
| Poverty* after initial hospitalization |  |  |  |  |  |  |
| OOP payment | All families<br>(n=315) |  | Government employed<br>(n=124) |  | Non-government employed<br>(n=191) |  |
|  | Impoverished | % | Impoverished | % | Impoverished | % |
| 1 month HHI | 65 | 21 | 6 | 5 | 59 | 31 |
| Cumulative** HHI | 29 | 9 | 1 | 1 | 28 | 15 |

HHI = household income \* Peninsular Malaysia HHI poverty line of MYR930 (2014) \*\* Cumulative by length of hospital stay

Overall 140 out of 370 families interviewed incurred CHE (38%). Malay families who were the majority had CHE incidence of 38%, Indians 30% and Chinese 20% while other minority ethnic groups had 50%. According to age group the highest incidence of CHE occurred among those less than 20 for both paternal (100%) and maternal (93%) categories. This was followed by the 20-29 age group (48% and 43% respectively). CHE mainly occurred in the lowest education group (85% and 75% respectively for paternal and maternal categories). CHE occurred mainly among fathers who were self-employed (52%) and in the private sector (39%) with lower incidence among government employed (13%). The highest incidence for CHE among mothers was in the home maker group (50%) followed by private sector (36%), self-employed (35%), and government sector (10%). CHE incidence followed a reducing pattern towards higher income quartiles with 76% of families in the first quartile affected compare to only 3% in the fourth quartile. More families from outside the district (49%) encountered CHE compared to families from within the district (22%).

Analysis of CHE among families by degree of prematurity revealed the highest incidence in the very preterm group (47%) followed by moderate preterm and extreme preterm groups (41% and 40% respectively) and to a lesser degree the late preterm group (30%). Analysis by preterm infant birth weight revealed the highest CHE incidence in the lowest birth weight category of less than 1kg (48%). CHE incidence fell as birth weight increased with the lowest incidence for those in the 2kg and more birth weight category (29%). 41% of families whose preterm infants were delivered normally had CHE compared to 36% for those whose preterm infants were born via Caesarean section. This was despite mothers who underwent Caesarean section having to stay in ward longer for postnatal care. CHE occurred most in families whose preterm infants had ward stay of 30 to 59 days (48%) or 60 days and more (44%). For preterm infants who stayed less than 30 days in ward the incidence of CHE among their families was 33%.

### Predictors of CHE

From univariable regression three clinical factors (LOS, gestation, birth weight) and five household factors (parental age, education and occupation, residential location and household income) were found to be significant. However multivariable regression showed only two factors to have significant association with the incidence of CHE namely residential location and household income (Table 5). The Hosmer–Lemeshow test showed goodness of fit (*p*=0.855). Families of preterm infants whose homes were located outside of the hospital district had increased probability of incurring CHE (OR = 2.855, p<0.05) while highest income quartile was protective against CHE (OR = 0.008, p<0.05). Clinical factors such as LOS, birth weight and gestation and household factors such as parental age, education and occupation were not significant factors for CHE.

**Table 5:** Predictors of catastrophic health expenditure (multivariable regression)

| Variable | OR | 95%CI | p |
| --- | --- | --- | --- |
| **Clinical factors** |  |  |  |
| Length of stay (< 1month = 1) | 1.960 | 0.962-3.993 | 0.064 |
| Gestation (34-36 weeks = 1) | 1.604 | 0.814-3.162 | 0.172 |
| Birth weight (<2kg = 1) | 1.777 | 0.839-3.761 | 0.133 |
| **Household factors** |  |  |  |
| Maternal age (<30 = 1) | 1.490 | 0.717-3.097 | 0.286 |
| Paternal age (<30 = 1) | 0.755 | 0.361-1.576 | 0.454 |
| Mother education (primary or lower = 1) | 0.893 | 0.255-3.118 | 0.859 |
| Father education (primary or lower = 1) | 2.302 | 0.689-7.693 | 0.175 |
| Mother occupation (homemaker = 1) | 0.734 | 0.373-1.444 | 0.370 |
| Father occupation (self-employed = 1) | 1.749 | 0.957-3.195 | 0.069 |
| Residential location (Out of hospital district = 1) | 2.855 | 1.583-5.148 | 0.000* |
| Household income quartile 2 (low)** | 0.288 | 0.138-0.602 | 0.001* |
| Household income quartile 3** | 0.089 | 0.038-0.212 | 0.000* |
| Household income quartile 4 (high)** | 0.008 | 0.002-0.034 | 0.000* |
| Log likelihood | 308.483 |  |  |
| $\chi^2$ (df) | 173.643 (13) | | |
| | $p = 0.000$ | | |
| Cox and Snell R square | 0.379 |  |  |
| Nagelkerke R square | 0.517 |  |  |
| Hosmer-Lemeshow test | $\chi^2$ (8)= 4.018 | | |
| | $p = 0.855$ | | |
| Observations | 370 |  |  |

\*Statistically significant at $p= 0.05$
\*\* Q1 is the reference group

## DISCUSSION

### Cost analysis

Total costs for government and non-government employed families of preterm infants were analyzed separately as government employed parents were entitled to full government subsidy of hospital cost. However, non-medical costs such as transport and food were borne by all families of preterm infants. It was observed that mean total cost was 18% higher for non-government employed families compared to government employed. Similarly the mean opportunity cost in the form of estimated income loss was 12% higher among them. Table 3 shows travel as the major cost contributor in both groups which correlates with results of previous studies where travel expenses accounted up to 60% of total cost during initial hospitalization (4, 5, 17). Travel cost for parents as they shuttle between home and NICU may be substantial especially if their preterm infants were referred to a distant regional centre. Other costs related to childcare at home and accommodation will likely follow suit (1).

### Financial sources

Approximately a third of families in this study had the benefit of full government subsidy to cope with hospital cost (direct medical cost). The remaining families who received partial government subsidy resorted to OOP payment for the remainder of hospital cost. Various strategies were used in this group but majority (one fourth) of them opted for current income of any household member followed by savings and borrowing from friends or relatives. No family in this study had the option of insurance coverage for hospitalization of their preterm infants. These findings are in line with the fact that for most of Asian population OOP payment is the primary method for health care financing (6, 7). A study in neighbouring Indonesia revealed borrowing as a dominant coping strategy among the poor to acquire health care during illness (18). In this study frequency of borrowing or selling assets for health care payment was lower than 26% noted in a study of lower and middle income countries (19).

Studies have found that majority of parents who visited their infants in NICU on a daily basis faced undisclosed financial difficulties mainly contributed by travel cost (20, 21). Furthermore they received little financial assistance due to limited and discretionary funds. A similar situation of undisclosed financial difficulties may exist in this study where more than two thirds of non-government employed parents perceived hospital cost payment as ‘not difficult at all’ or ‘not that difficult’ and less than a third found it to be ‘difficult’ or ‘very difficult’. This was despite the fact that more than a fifth of them were living below the poverty line, more than half suffered income loss and close to a fifth were unable to make full payment of the partially subsidised amount upon discharge. Despite full government subsidy of hospital cost government employed parents received full too experienced income loss, impoverishment and CHE but at a comparatively much lower rate than their counterparts.

Overall, 15% of families this study were living below the income poverty line and the overwhelming majority was from the non-government employed group. This was surprisingly very much higher than the 0.6% incidence of poverty for Malaysia and 0.3% for the state of Kedah in 2014 (14). With exclusion of families below poverty line further analysis was done and it was found that up to a fifth of families above poverty line suffered impoverishment when one month of HHI was used for initial hospitalization cost but this improved to less than a tenth when cumulative HHI (according to LOS) was used. Subsequently analysis was performed on government and non-government employment status to assess the impact of full and partial subsidy of hospital cost on impoverishment. Up to a fifth of non-government employed families were already living below the income poverty line prior to initial hospitalization compared to only 2% of government employed. Following exclusion of families below poverty line further scrutiny found that with a month’s HHI used for initial hospitalization cost, close to a third of non-government employed families became impoverished compared to just 5% of government employed families. When cumulative HHI (comparable to LOS) was used the situation improved to 15% in non-government employed families and just 1 affected government employed family. It appeared that government employed families were better insulated against poverty and impoverishment following OOP payment for health care. Due to an array of employment benefits civil servants in Malaysia may be less susceptible compared to those in private sector or self-employed. They enjoy job security, fixed working hours, regular income, wage hikes and higher minimum wage rate. Added perks include fully subsidized healthcare, low interest government loans, paid leave and a lifelong pension scheme that they can rely upon for financial stability.

In this study families in lower income quartiles were more likely to incur CHE compared to those with higher income which was consistent with previous studies (22–26). On a global scale, an income and expenditure survey data for 60 countries revealed household income as among the important determinants of CHE (27). The burden of healthcare payment for those with higher income become proportionately smaller due to their higher capacity to pay. In contrast, households with low income have very little capacity to accommodate health care spending without compromising daily basic needs (28). Consistent to the findings that incidence of CHE has a direct relationship with distance from healthcare facilities where CHE incidence was lower with reduced distance, in this study those families who resided at rural areas had more expenditure for travelling to the hospitals which were located in town (29). WHO reported in a regional study on health financing that distance to the health facility is a key determinant for access and inadequate access to the health facility is a determinant for CHE (30). Rural households have higher risk of CHE because of lower socio-economic status and longer traveling distance to healthcare facilities compared to those residing in the urban areas (29, 31–33).

In this study residential location was one of the predictive factors for CHE with travel expenditure as the main cost contributor for families of preterm infants during initial hospitalization. Thus higher travel expenditure may be a likely reason why families staying outside of the hospital district had higher probability of experiencing CHE. Considering these findings, rooming-in service for mothers at NICUs should be improved as it can reduce the frequency of travelling for visitations and associated costs. Rooming-in for mothers is an excellent alternative as it benefits both mother and baby such as encouraging breastfeeding, skin to skin contact and promotion of growth (34–37). Another measure that can be implemented to reduce family financial burden is complementary financial assistance as HHI was found to be a predictive factor for CHE in this study. Furthermore it was demonstrated that despite universal health coverage (UHC) through extensive government subsidy for hospital cost the incidence of CHE and impoverishment among families of preterm infants was still high and it affected non-government employed families more. This was contributed by OOP spending for direct non-medical costs and losses from opportunity costs. Government subsidy for hospital fee only addressed direct medical cost when direct non-medical cost such as for travel was the cost driver for families during initial hospitalization. The effect was especially pronounced among parents who were not government employed who received partial subsidy of hospital fee compared to those government employed who received full subsidy. Families in need of financial support were referred to the hospital social support department but it has restricted ability to assist due to limited annual financial allocation. Studies have revealed that only a quarter of patients received financial help of any source while the rest had little or no help from relevant authorities (17, 38). Personnel from NICU and hospital social support must identify families in need and the hospital social support department should be strengthened with adequate funds. Complementary financial assistance will help alleviate the financial burden during initial hospitalization and allow families of preterm infants to provide for subsequent growth and development needs. To enhance UHC, affordable pre-payment for infant healthcare should also be considered by the government. This is especially for those who are not government employed and not entitled to full subsidy of hospital services. A social health insurance service that provides coverage during pregnancy and the infant’s early years should be implemented to reduce OOP payment. A comprehensive coverage would include pregnancy complications, initial hospitalization cost such as NICU admission, treatment for congenital conditions and management of growth and developmental disorders associated with prematurity that may present later.

Malaysia has been identified alongside Thailand and Sri Lanka as low to middle income countries that have managed to curb both the OOP share of health financing and the catastrophic impact of direct payments through UHC (15). A government (tax-based) subsidised public health care coupled with private sector health care services has produced a progressive health financing system.

According to the Ministry of Finance economic report 2017 the government subsidy was more than 23 billion Malaysian Ringgit in 2017 across public sectors (39). The health subsidy in Malaysia is at more than 90% meaning more than 90% per cent of the health costs for every patient is borne by the government. Despite UHC and heavily subsidized direct medical (hospital) cost for families of preterm infants, findings in this study suggest that it was inadequate to shield them from CHE and impoverishment. This can be attributed to OOP payment for direct non-medical costs (travel, food, accommodation, child care) and losses incurred in the form of indirect (opportunity) cost. The impact was more pronounced upon those who were not government employed.

There were limitations in this study. Recall bias was a possibility for monthly expenditure and income loss estimates during interview but usually important incidents such as this would not be easily forgotten by the family. The 2014 average income poverty line used was for Peninsular Malaysia in general and did not discern between urban and rural poverty lines as family residence data was not specified for urban or rural locations. Since the Peninsular Malaysia average poverty line used in this study was higher than the rural poverty line it possibly yielded more conservative rates for poverty and impoverishment. However its strength lies in the multiple measures both objective and subjective were utilized to illustrate the gravity of problem faced. They included ability, perceived difficulty and coping strategies to settle hospital cost, income loss suffered, CHE and impoverishment. Sub-analysis of these measures between government and non-government employed families exposed the latter as among sections of the population (private, self and non-employed) vulnerable against financial burden of healthcare despite UHC in this country. This study was the first locally or abroad to determine the incidence and predictive factors for financial burden among families of preterm infants during initial hospitalization.

To obtain better understanding about perceived difficulty of OOP payment for healthcare among parents of preterm infants a qualitative study should be conducted. In-depth study will provide better insight into the multitude of factors that influence the perception and ultimately ability to pay. This study looked at incidence of impoverishment but not the effect over time. The effect of impoverishment upon families of preterm infants should be followed up to determine the true impact, whether transient or sustained. Following discharge from initial hospitalization complications such as chronic lung disease, growth and development problems and learning disabilities require rehospitalisation, outpatient medical care, early intervention and special education. These necessitate assessment of care provider, family and societal costs beyond the neonatal period as it has been estimated to reached billions of dollars annually with more than three quarters of the cost delivered during early childhood (0-5 years) (2). Coupled with findings from this study it would provide a broader spectrum of the economic costs of preterm birth in Malaysia.

## Acknowledgements

The authors would like to acknowledge consultant pediatricians Dr. Thiyagar Nadarajaw, Dr. Tan Ying Beih and Dr. Choo Chong Ming for their willingness to participate in this study. Special mention to NICU doctors and nurses of both study centers who diligently assisted in data collection and analysis. We would also like to express our gratitude to the Director General of Health, Malaysia for permission to conduct and publish this study.

## Author contributions

Conceptualization – Hadzri Zainal, Maznah Dahlui, Tin Tin Su

Data Curation – Hadzri Zainal

Formal Analysis – Hadzri Zainal, Maznah Dahlui, Tin Tin Su

Funding Acquisition – Hadzri Zainal, Maznah Dahlui, Tin Tin Su

Investigation – Hadzri Zainal

Methodology – Hadzri Zainal, Maznah Dahlui, Tin Tin Su,

Project Administration – Hadzri Zainal

Resources – Hadzri Zainal Software – Hadzri Zainal

Supervision – Maznah Dahlui, Tin Tin Su

Validation – Hadzri Zainal, Maznah Dahlui, Tin Tin Su

Visualization – Hadzri Zainal

Writing – Original Draft: Hadzri Zainal

Writing – Review & Editing: Hadzri Zainal, Maznah Dahlui, Tin Tin Su

